# A phrenic-sparing cephalic reanastomosis model: Acute effects and implications

**DOI:** 10.1101/2025.05.06.652103

**Authors:** Michael Lebenstein-Gumovski, Alexander Zharchenko, Tanzila Rasueva, Eugene Sobol, Pavel Petrov, Kirill Eskov

## Abstract

**Background:** Head transplantation (HT), also known as cephalosomatic anastomosis (CSA), is a surgical procedure proposed as a potential method to extend lifespan in cases of terminal bodily failure. CSA requires a thorough evaluation of each step using appropriate experimental animal models, with pigs being particularly suitable due to their anatomical and physiological similarities to humans. The critical challenge in HT is spinal cord fusion, which current research suggests can be facilitated using fusogens. To refine the technical aspects of the procedure, we conducted head replantation in a pig model using polyethylene glycol (PEG)-chitosan conjugate (Neuro-PEG). In this article, we evaluate the technical aspects of this procedure, the rate of recovery of ventilation after spinal cord fusion, and phrenic sparing in a model of autologous decapitation-reanastomosis in a single swine.

**Methods:** A Hungarian Mangalica pig was submitted to surgical separation of the head while maintaining blood flow to the brain through cannulation of the primary cervical vessels. After cephalic separation, the head was reconnected, and the cervical spinal cord fused.

**Results:** The animal was weaned from ventilation after 5 h and kept on spontaneous breathing. The animal regained full consciousness, demonstrated early signs of sensory recovery, and restored brain functions.

**Conclusion:** Pending further confirmatory studies, a phrenic-sparing cephalic reanastomosis with spinal cord fusion using fusogens has clarified the technical aspects of the procedure. A head replantation model with complete vascular cannulation was developed, resulting in the recovery of spontaneous breathing and brain functions.

## INRODUCTION

Cephalosomatic anastomosis (CSA), a.ka. Head transplantation (HT) traces its origin to the early XX century and built up momentum in the 1950s–1970s with the work of Vladimir Demikhov^[9]^ in the Soviet Union and White^[32–34]^ in the USA: these early pioneers transplanted the heads of dogs andmonkeys and kept them alive for a few days.^[23]^ However, technical issues (e.g., reconnection of the spinal cord and lack of immunosuppressants) and ethical concerns (the Ick Factor) cast a pall on the whole undertaking, and the field fell into disgrace.

The primary challenge remains the inability to restore spinal cord function following its connection. This is attributed to the dominance of destructive processes associated with secondary spinal cord damage over its regenerative capacity. The discovery of the mechanism of fusogenic repair has provided a fresh perspective on this longstanding issue. Fusogens initiate axonal fusion before secondary damage occurs, enabling neurophysiological responses within minutes of application. The first promising results with fusogens were achieved in the field of peripheral nerve surgery.^[2,3,13,25,26]^ Detailed development of protocols and theoretical substantiation for the feasibility of CSA was undertaken by Sergio Canavero in 2013, culminating in the presentation of his GEMINI project.^[4–7]^ Since then, theoretical mechanisms of spinal cord fusion have been explored, and various fusogens have been studied in experimental models of spinal cord injury.^[20,35]^ Some of the most promising fusogen combinations (Polyethylene glycol [PEG]-chitosan, Neuro-PEG) have demonstrated the ability to restore spinal cord function within a few days following complete transection in experiments on large animal models.^[18,19,21]^ CSA experiments have primarily been conducted on small animal models.^[16,23]^ However, to date, CSA, in large, highly translational animal models, remains a significant technical challenge.

The relaunch of a head transplant initiative (HEAVEN) early in 2013^[6]^ culminated in a full human cadaveric rehearsal in 2017,^[7]^ along with the demonstration that a severed spinal cord in this setting can be functionally restored.^[15–29]^

One key aspect of the procedure is the quick return of spontaneous ventilation. As is well known, prolonged mechanical ventilation comes with an increase in mortality, along with muscle wasting (which would hinder post CSA rehabilitation), pressure ulcers, infections, pulmonary embolism, and others.^[17,22,31]^ In the 2017 paper, the HEAVEN authors wrote:

“*… Two options were considered: transecting the phrenic nerve and then reanastomosing/fusing it during the final stages of the cephalic anastomosis or simply sparing it. These possibilities would also mean transecting the spinal cord at different levels: between C2 and C3 myelomeres or C5 and C6 myelomeres. Based on our success in reestablishing electrical continuity of the spinal cord early on within 4 weeks with behavioral recovery (…) and much less experience with accelerated, fusogen-supported peripheral nerve reconstruction/fusion (…) in addition to the need to maintain breathing once the chimera is reawakened, we reached the compromise of saving the C4 and C5 components and sacrificing the C3 component. Thus, the vertebral column was transected at the C3-C4 level; we expect that this construct will be sufficient to support spontaneous respiration after the restoration of electrical continuity of the spinal cord. In any case, this is being tested in an animal model*.”^[28]^

To determine the technical feasibility of such a procedure for further experiments, we developed a swine head replantation model. In this preliminary study, we set out to test the effects of cephalic re-anastomosis in a swine model, specifically the postoperative recovery of spontaneous ventilation following complete spinal cord transection at C4 and subsequent spinal cord fusion, as well as the impact of the vascular cannulation procedure on the primary functions of the brain.

The present model involving autologous cephalic transection and reconnection is ideal, as no immunosuppression is required and neuroprotection is unnecessary as the flow is guaranteed during the entire procedure (see ahead).

## MATERIALS AND METHODS

A Hungarian Mangalica pig (female, m = 25.0 kg. *n* = 1), treated in accordance with the European Convention for the Protection of Vertebrate Animals used for Experimental and Other Scientific Purposes (European Treaty Series - No. 123, March 18, 1986), was submitted to the experimental procedure. The Local Ethics Committee approved all actions. The preoperative neurological examination was normal, including papillary, corneal, and pharyngeal reflexes, eye movements, and sensory and motor (Individual Limb Motor Scale) scores.

For ethical reasons and to avoid any undue protracted pain, it was decided to assess the results over 12 h postoperatively and then euthanize the animal.

In the days preceding surgery, we prepared neuroPEG, that is, chitosan with a molecular weight of 15 kDa (Merck, Germany), stabilized by photo-crosslinking, in a chemical conjugate with PEG-600 (Merck, Germany), at a concentration of 20 mg/mL. The resulting mixture was sterilized and stored in sealed tubes at a temperature of 4°C (Neuro-PEG).^[18,19]^

### Surgery

General anesthesia was obtained with a mixture of IM tiletamine/zolazepam (13.2 mg/kg, Zoletil, Virbac, France) and Xylazine (0.15 mL/kg, Alfasan Int., Netherlands) followed by propofol (3 mg/kg, Propofol-Kabi - Fresenius Kabi, Germany) through a central venous catheter. The animal was curarized (Rocuronium, 0.4 mg/kg, Kabi-Fresenius Kabi, Germany), and a urinary catheter was inserted. A low tracheostomy was fashioned, and a long endotracheal tube was inserted; the animal was ventilated with an O_2_ (24%)-air mixture.

### Surgery: Anterior approach

The animal was placed supine and secured to the operating table. The head was held in place with a custom-made fixation device, with the neck in a neutral position. The upper limbs were retracted to provide better access to the anterior surface of the neck. A wedge-shaped incision was made as low as possible on the anterior surface of the neck [Figure 1]. The skin was lifted from the muscle layer and rolled outward both cranially and caudally; stay sutures were applied [Figure 2]. The superficial and deep muscles of the anterior surface of the neck were carefully isolated, dissected at their bony base as low as possible, ligated, and thrown back cranially [Figure 3]. The thyroid gland and trachea were exposed and the latter transected below the thyroid gland and shifted cranially, with the distal portion intubated. The recurrent laryngeal nerves were visualized and deliberately transected to facilitate the procedure. The esophagus was transected inferiorly, and the stumps moved laterally. The cervical neurovascular bundles were isolated and pushed aside. The spine was accessed at C3-C4-C5. The prevertebral fascia and anterior longitudinal ligament were dissected, and a Caspar Cervical Distractor (Aesculap, Germany) was applied at C3 and C5. After moderate distraction, a corpectomy of C4 was performed [Figure 4]. The posterior longitudinal ligament was visualized and dissected, and the anterior surface of the dural sac was visualized. At this point, the carotid arteries and the jugular veins were sequentially transected, cannulated, and heparinized [Figure 5]. Finally, the ends of both vagi were soaked in a cold 2 mM chlorpromazine solution (Pharmacel, Russia) and transected.

**Figure 1:**
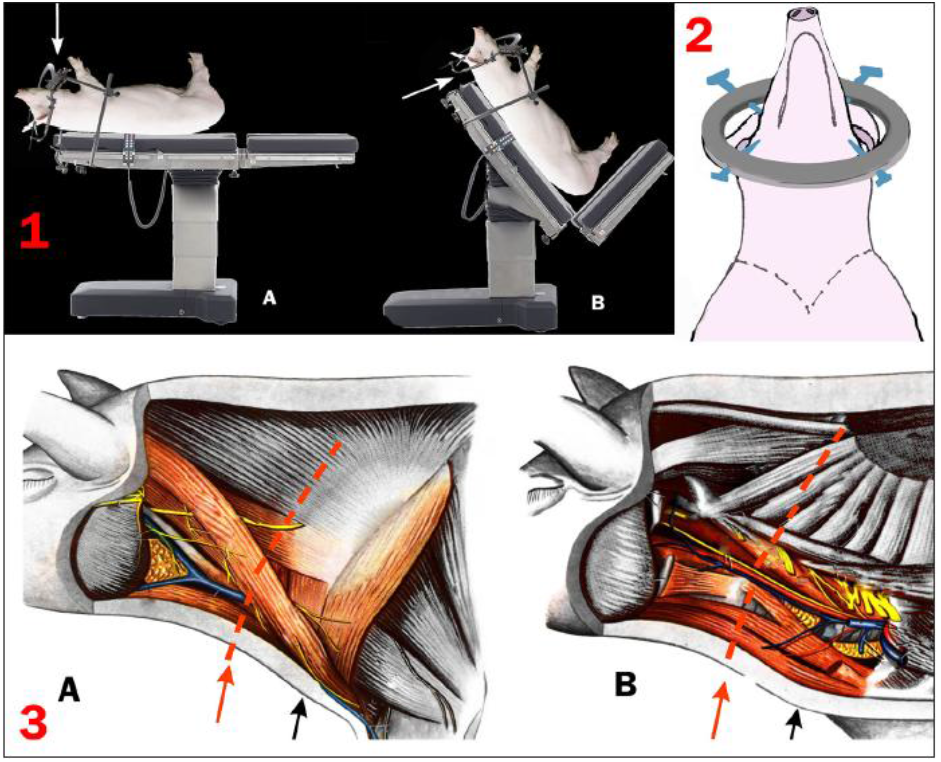
1: Position of the animal for the initial anterior approach (a) and position of the animal for the later posterior approach (b) (arrows: angles of surgical attack); 2: layout of the position of the animal’s head inside the clamp and skin incision for the anterior approach (dotted line); 3: muscles and neurovascular bundles of the pig neck: (a) Superficial muscles of the neck; and (b) deep muscles of the neck. The structures involved in the anterior approach are highlighted in color. Black arrow: Projection of the skin incision and muscle separation; Red arrow: Projection of the level of the spinal section (adapted from: Popesko, P. [Atlas of Topographical Anatomy of the Domestic Animals: Voulme I, Head and Neck], Bratislava: Priroda, 1978. - 211 p. In Russian).

**Figure 2:**
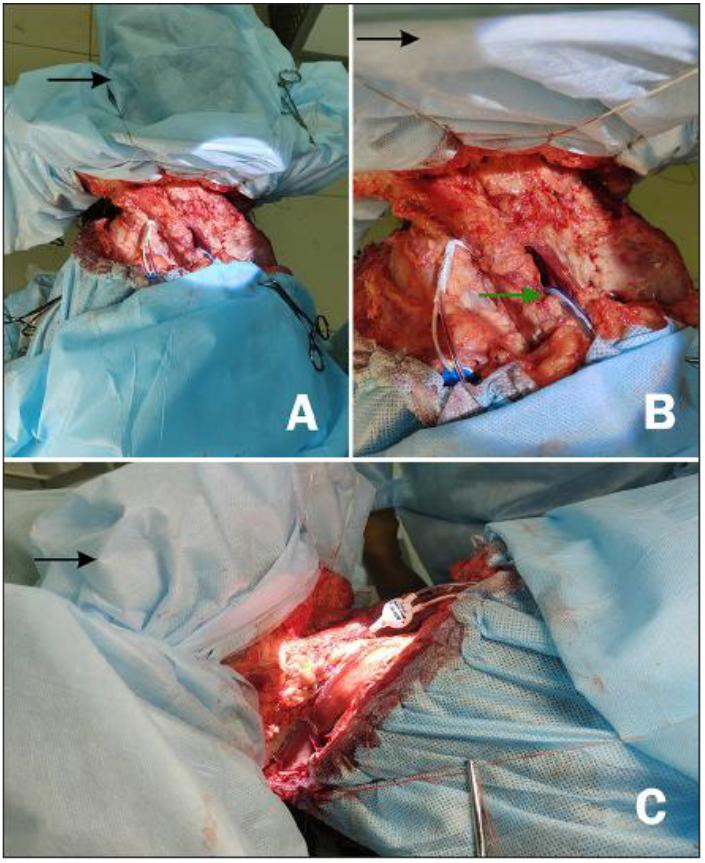
Anterior approach after separation of the skin flaps. (a) General front view. (b) Front view, close-up. Arrow: Endotracheal tube. (c) Side view of the surgical approach. Blackarrow: Cranial end.

**Figure 3:**
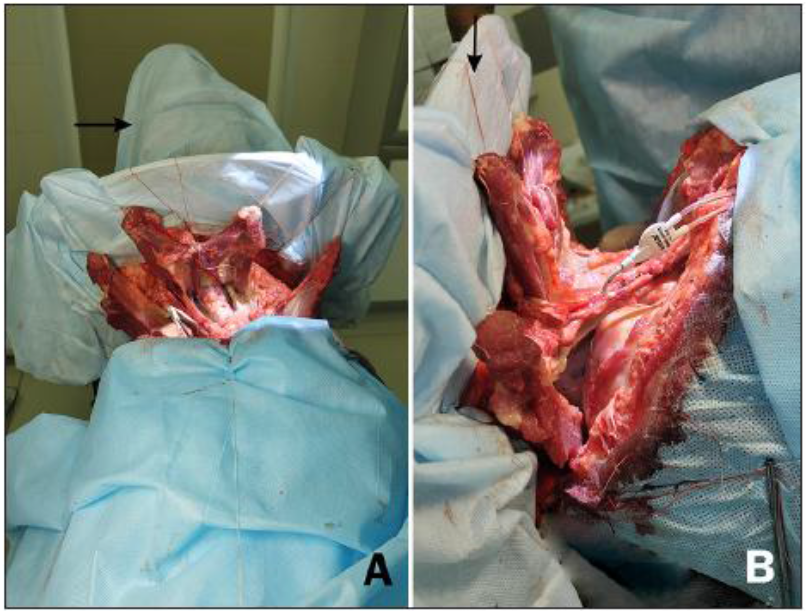
Anterior approach. The neck muscles are separated and elevated with ligatures. The trachea is visible. Black arrow: Cranial end. (a) Anterior view and (b) lateral view.

**Figure 4:**
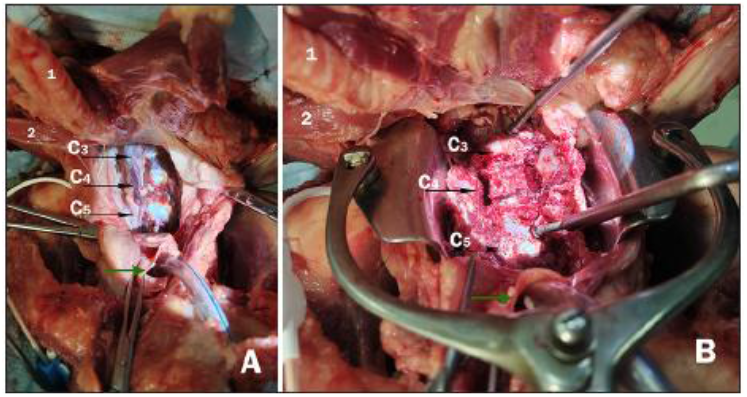
Anterior approach. (a) Approach to the anterior surface of the bodies of the C3–C5 vertebrae (black arrows). (b) A Caspar distractor is installed in the C3 and C5 vertebral bodies. Initial C4 corpectomy. The cranial end of the trachea (1) and the cranial end of the esophagus (2) are visible. Green arrow: Distal end of the trachea with the endotracheal tube.

**Figure 5:**
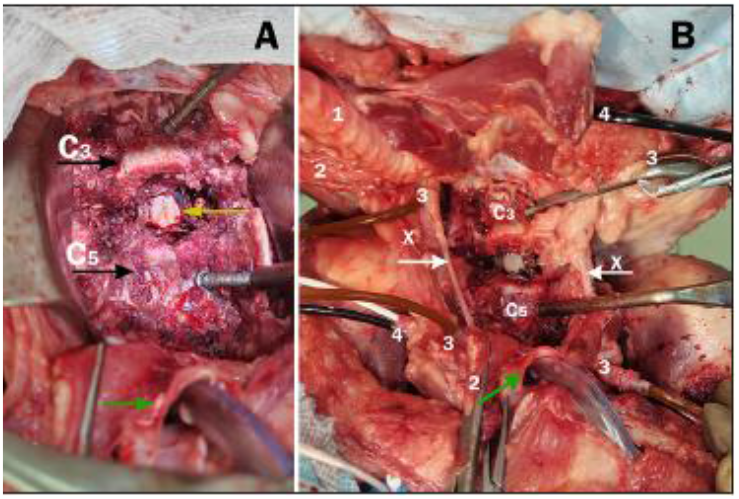
Anterior approach. A corpectomy of C4 is performed. (a) Black arrows: bodies of the C3 and C5 vertebrae. Yellow arrow: dissected posterior longitudinal ligament and anterior surface of the dural sac. (b) Bird’s eye view of the approach at this stage. The trachea (1), cranial and caudal ends of the esophagus (2), cannulated proximal and distal portions of the carotid arteries (3), cannulated jugular veins (4), and the bodies of the C3 and C5 vertebrae are visible. White arrows: vagi; Green arrow: trachea with the endotracheal tube. The cranial end is topmost.

### Posterior approach and separation

Subsequently, the operating table was moved to a semi-vertical position [Figure 1] to provide access to the back of the neck.

The initial cutaneous incision was extended semicircularly, with additional longitudinal incisions overlying the spinous processes. The skin was dissected, folded cranially, and ligated. The posterior group of superficial muscles of the neck was partially dissected. The spinous processes of the C3-C4-C5 vertebrae were dissected free, and a retractor was applied [Figure 6]. Thereafter, all three processes were resected, and the arch and remnants of the pedicles of the C4 vertebra were completely removed.

**Figure 6:**
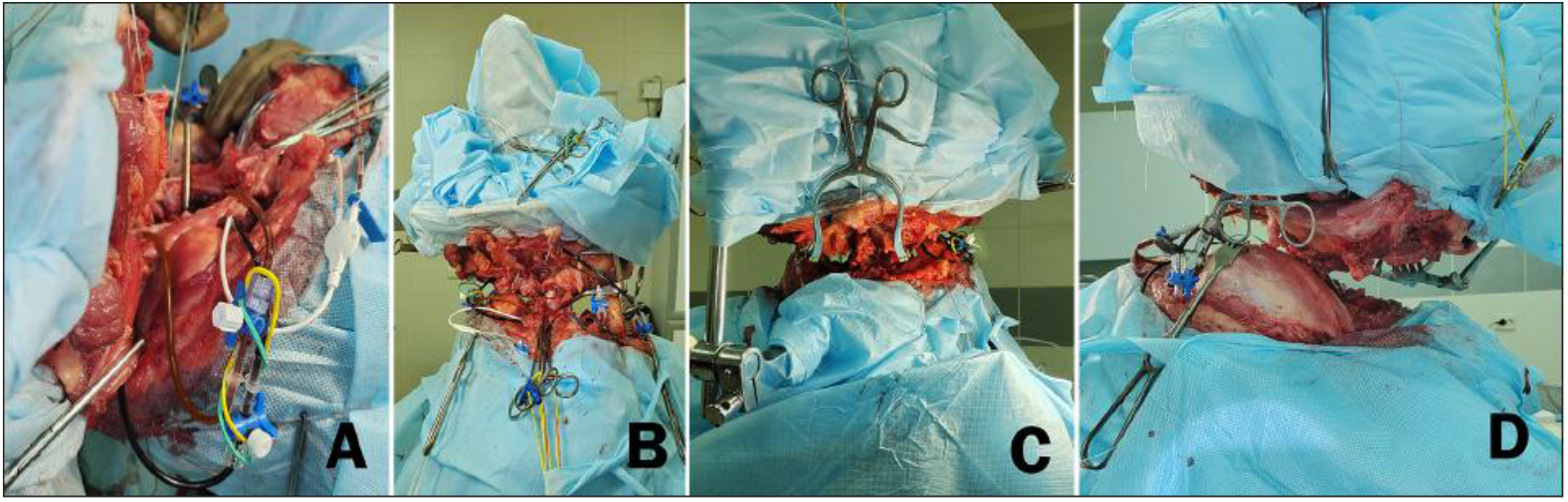
Intermediate stage with completed anterior and posterior approaches. (a) Side view, right: Cannulated vessels are visible. The cephalic end is on the left. (b) Anterior view. The head is held in the fixation ring and is completely separated, except for the spinal cord. (c) Posterior view. The cephalic end is above. (d) Lateral view left (head: topmost).

One vertebral artery was spared to ensure blood flow to the posterior circulation, while the other was sectioned.

At this point, the head was almost completely separated from the body, except for the spinal cord [Figure 6].

A titanium mesh implant (Medbiotech, Belarus) was installed at C4: screws and longitudinal rods (LLC Osteomed, Russia) were placed transpedicular at C3 and C5 for posterior fixation [Figure 7]. At this stage, circular sleeves of artificial dura mater (MedPrin Biotech GmbH, Germany) were applied to the dural sac using cyanoacrylate glue (BBraun, Germany) as part of the 360° dura sealing method that we propose.

**Figure 7:**
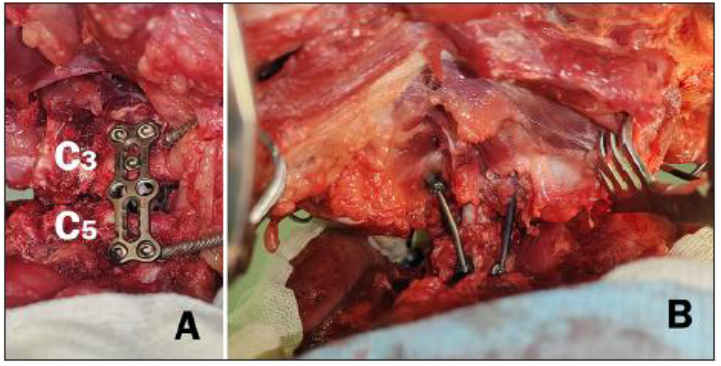
(a) View from the anterior approach. Anterior cervical plate and interbody cage. The distractor has not yet been removed. (b) View from the posterior approach. Transpedicular fixation. (Head: topmost).

Then, 12 stay sutures (Prolene 6/0, Ethicon, Johnson and Johnson, USA) were applied, and the dura was covered with ice slurry. Thereafter, the dura mater was circularly incised and retracted [Figure 8].

**Figure 8:**
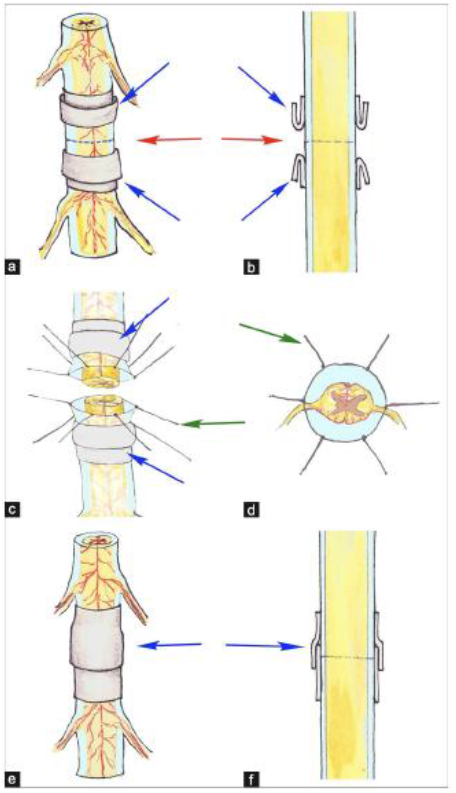
Scheme of circular 360-sealing of the dura mater using sleeves. (a and b) Installation of sleeves made of artificial dura mater at a distance from the incision. The edges of the sleeves are everted (blue arrows). The red arrows and dotted lines show the area of the proposed incision. (c and d) Circular incision of the dura mater and dissection of the spinal cord. The edges of the membrane are sutured with ligatures (green arrows); blue arrows show the sleeves. (e and f) After approximating the ends of the spinal cord and tightening the ligatures, the sleeves are repositioned, and the sleeve is fixed with glue (blue arrows).

Finally, the spinal cord was completely transected at C4.

### Reanastomosis

The C3 and C5 bodies were released from the distractor. Posteriorly, the two stumps of the cord were shortened a few millimeters and then fused by applying 5 mL neuroPEG. The 12 ligatures that held the dura mater were pulled together and knots fashioned. The artificial dural sleeves were pulled over each other in an overlapping fashion, completely sealing the dural sac defect [Figure 8]. The overlap area was then glued with cyanoacrylate glue (BBraun, Germany). The posterior transpedicular fixation system (LLC Osteomed, Russia) was secured. Anteriorly, a C3-5 cervical plate (LLC Conmet, Russia) was installed [Figure 7]. Finally, all previously sectioned structures were sutured back: esophagus, trachea, and muscles. The vagi were sutured and, after slight shortening to ensure a fresh fusion surface, undiluted PEG 600 (1 mL) was applied.

Finally, the animal was then intubated with an orotracheal tube (N 6.5, Alba Healthcare LLC, USA), and the tracheostomy window closed. Standard closure of the skin followed. The Central Venous Access and wound drainage were left *in situ*.

Total operation time was 17.5 h from the first incision and 19 h in total.

Postoperatively, painkillers were administered according to international guidelines,^[1]^ anticoagulation was instituted,^[14]^ antibiotics were infused, and the animal was kept in a special isolated box. Parenteral nutrition and hydration were provided as per guidelines.^[1]^

## RESULTS

The animal was kept sedated during the first 5 h after the operation and artificially ventilated. Then, the animal was awakened and allowed to breathe spontaneously [Figure 9a].

**Figure 9:**
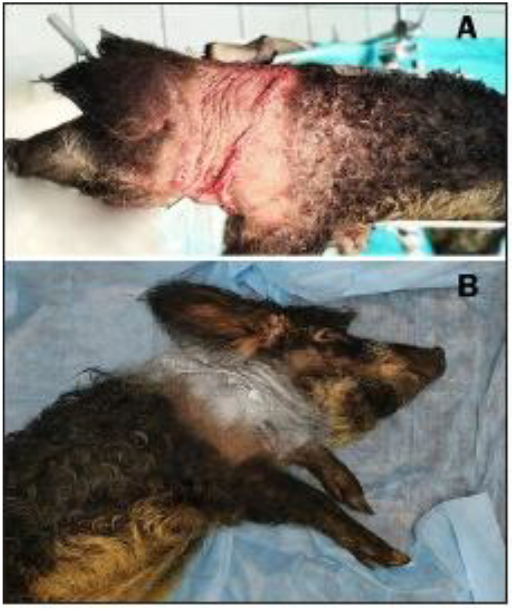
Animal after head replantation. (a) 5 h after surgery. Residual sedation is maintained, and the animal is transferred to spontaneous breathing. (b) Animal 10 h after surgery [Video].

Breathing was weak and intermittent, and mask oxygenation was periodically required during the 1^st^ h.

Pupillary, corneal, and swallowing reflexes were normal, and no sign of ophthalmoplegia was seen. Moderate miosis was observed due to loss of the sympathetic drive. The animal was fully responsive to the surrounding environment, including sounds, and willingly drank water from a syringe. Muscle tremors and attempts at muscle contraction to pain stimulation were observed in the limbs. Nociception was assessed with a pin at three control points on each limb, as well as on the back and the anterior surface of the trunk. When all points were stimulated, the animal responded with a squeal, but no active movements were recorded.

Ten hours after the operation, the picture remained unchanged [Figure 9b and Video 1]. As per protocol, the animal was euthanized at 12 h to study the acute effects of spinal cord fusion with histological analysis (*unpublished observations*).

**Video 1:**
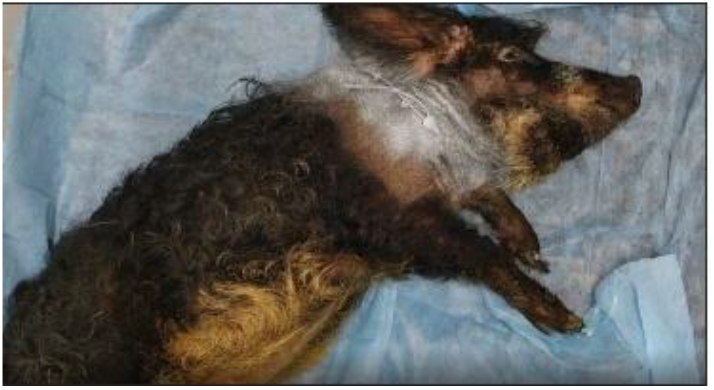
Animal 10 hours after surgery.

## DISCUSSION

Swine are increasingly studied as animal models of human disease. The anatomy, size, longevity, physiology, immune system, and metabolism of swine are more like humans than traditional rodent models. In addition, the size of swine is preferred for surgical placement and testing ofmedical devices destined for humans. These features make swine useful for biomedical, pharmacological, and toxicological research.^[24]^ This operation holds significant importance for understanding the technical and anatomical-physiological aspects of HT in a large animal model.

The described procedure integrates both the explanation and separation of the head from the donor and recipient, as well as the creation of CSA. Operation duration was recorded, key procedural sequences were evaluated, and adjustments were made to the initial algorithm.

In addition, we tested the method of circular plastic surgery and dura mater sealing. Fusogenic fusion was applied to both the vagus nerve and spinal cord. The observation of early sensory responses aligns with previously obtained data.^[18,19]^

There are also differences and we highlight two of special interest to the present work. In pigs, the phrenic nerves are formed by the ventral branches of C5, C6, and C7 spinal nerves, with C5 and C7 roots particularly thin. These roots converge in a single trunk at the level of the C7 vertebra and, then, are distributed into the diaphragm muscle.^[10]^

The pig has a large fine-caliber vessel plexus providing blood to the neck area, from which flow reaches both the spinal cord and the base of the brain. This rete mirabile (RM) is supplied by bilateral ascending pharyngeal arteries. Bilateral anterior cerebral arteries, middle cerebral arteries, and the basilar system lie rostral to the RM.^[11]^ Furthermore, large bilateral vertebral arteries feed the circle of Willis in pigs, with abundant small branches. The first two branches on each side are major tributaries, which arise in a right-angle fashion from the vertebral artery and head toward the spinal cord in the cervical area: they are of major importance for the spinal cord above the segmental arteries.^[30]^

In this work, we show that very early weaning from post-cephalic reanastomosis mechanical ventilation is feasible, although not optimal, as the animal had to be assisted with supplemental oxygen. The rationale for the chosen level of spinal cord transection has been previously established.^[4–7]^ Due to the rapid degeneration of neural tissue, it is essential to prepare the recipient and donor spinal cord stumps much longer than would be required for alignment and re-cut them immediately before fusion. Neurologically, it is justified to preserve most of the cervical enlargement in the donor body, as this region contains the origin of the phrenic nerve and innervation for the upper limbs. Therefore, spinal cord transection at the C3 segment is deemed appropriate.^[4,6]^ Yet, it is clear that the concept put forth by previous authors Ren *et al*.^[28]^ as outlined in the introduction, is vindicated. The mere acute application of fusogens leads to a very quick recovery of electrical conduction across the sectioned cervical cord. Previous studies demonstrated that animals so treated – from rodents to primates – show the first visible signs of motor recovery at 48 h,^[15,18,19,29]^ but electrophysiologically motor and somatosensory-evoked potentials display initial recovery of conduction within the hour.^[29]^

The vagus nerves play a critical role in systemic physiological regulation, making their rapid restoration essential. Fusogenic nerve repair enables the recovery of action potentials within minutes following neurorrhaphy, offering a promising approach for accelerated functional recovery. In this context, peripheral nerve repair should incorporate the protocols of De Medinaceli^[8]^ and Bittner *et al*.,^[2,3]^ as well as strategies involvingcalcium chelators^[27]^ to optimize outcomes. It is worth noting that the organs – the heart and kidneys, after transplantation, do not have parasympathetic innervation, and the absence of vagal innervation does not affect their functions. Similarly, relatively normal intestinal motility is restored within a month after isogenic intestinal transplantation.^[4]^

When preparing the recipient’s head, preservation of the recurrent laryngeal nerves is essential. In this experiment, we did not aim to maintain their arch under the aorta and brachiocephalic trunk; however, this step will be necessary for a complete transplant.^[4]^

In addition, damage to the sympathetic trunk may result in Horner’s syndrome and other autonomic disturbances in the recipient, leaving the question of its reconstruction an open area for further study.

However, challenges persist regarding the vertebrobasilar system, brainstem, and especially the recipient’s spinal cord. The brainstem, cerebellum, parts of the occipital and temporal lobes, as well as basal nuclei, receive blood supply from the vertebral artery system. Even with well-developed collateral circulation and a functional circle of Willis, the carotid system alone cannot fully compensate for vertebrobasilar insufficiency. The entire cervical spinal cord is vascularized by radiculomedullary arteries originating from the vertebral artery system, which, in turn, branches from the subclavian arteries. During head severance, the vertebral arteries are transected, disrupting the blood supply to the spinal cord. To address this, bilateral shunts between the external carotid artery and the ipsilateral vertebral artery above the transection site in the recipient must be established to restore adequate blood flow.^[3]^

This revascularization should maintain blood flow both above and below the anastomosis, ensuring adequate perfusion of the recipient’s spinal cord. An alternative approach involves creating a high-flow venous shunt between the subclavian artery and the V3 or V4 segment of the vertebral artery on each side. Such a shunt could be established in the recipient 2–3 weeks before transplantation and subsequently cannulated at the time of head separation to preserve blood circulation. The effectiveness of descending revascularization of the spinal cord by such methods, as well as the sufficiency of ascending revascularization of the vertebrobasilar basin, remains a subject of discussion and further research.

Current research on fusogenic spinal cord repair primarily focuses on the use of PEG and its derivatives.

Previous work showed that the experimental species plays a role in the rate of recovery after spinal cord fusion: rodents recover from paralysis within days, dogs within weeks, and primates within months.^[15,29]^ In our studies exploiting a combination of fusogens, final recovery in swine was similar to canines,^[19]^ although arguably faster. We plan to compare our fusogenic combination^[19]^ with the only published PEG study in monkeys^[29]^ in a head-to-head study. We incidentally notice how Bittner’s protocol of peripheral nerve fusion^[2,3]^ has never been tested for spinal cord fusion (see commentary to ref 20). We also see a potential adaptation of the De Medinaceli approach to neural fusion^[8]^ in the present context.

Despite a number of significant differences and aspects that must be addressed in a complete transplant, the primary issues still under discussion involve the preservation and restoration of neural structures, ensuring stable perfusion of the brain and spinal cord, and preventing ascending spinal cord edema.

## CONCLUSION

We developed a swine model of HT that can be leveraged to explore several aspects of the technology involved in HT.

## Acknowledgments

We thank Andrey Panferov, Alina Nurislamova and the editors for language support of this article.

## Ethical approval

The research/study approved by the Institutional Review Board at Stavropol State Medical University, number 97, dated April 15, 2021.

### Declaration of patient consent

Patient’s consent was not required as there are no patients in this study.

### Financial support and sponsorship

Nil.

### Use of artificial intelligence (AI)-assisted technology for manuscript preparation

The authors confirm that there was no use of artificial intelligence (AI)-assisted technology for assisting in the writing or editing of the manuscript and no images were manipulated using AI.

## How to cite this article

Lebenstein-Gumovski M, Zharchenko A, Rasueva T, Sobol E, Petrov P, Eskov K. A phrenic-sparing cephalic reanastomosis model: Acute effects and implications. doi:

## COMMENTARY

In this paper, these Russian neurosurgeons examine a key point of a full head transplant, that is, the recovery of spontaneous breathing. After having confirmed in a swine model that spinal cord fusion is indeed possible (their refs 8-9), they moved on to a swine model of head transplantation they developed. This model is commendable for its simplicity: no need to sacrifice two animals, as one is sufficient; no need for immunosuppression, as the entire surgery is autologous; and finally, no need for neuroprotection as the head is never deprived of circulation. As such, this model is ideally suited to test the recovery of sensory, motor function, and vegetative functions after spinal cord fusion. In this case, the focus of the authors has been on the recovery of spontaneous breathing. And indeed, the animal recovered this function. Actually, the full GEMINI spinal cord fusion protocol involves spinal cord electrical stimulation at the level of fusion, and we already know that this boosts the mere application of fusogens (*unpublished observations*). Yet, this is an extraordinary result for the Russian team.

Clearly, what was (and still is in some quarters) considered impossible is actually quite feasible. As the father of astronautics, Russian Constantin Tsiolkovsky, said: *The impossible today will be possible tomorrow*. It seems that tomorrow is, in fact, today.

Sergio Canavero, MD (US FMGEMS)

HEAVEN/GEMINI Global Initiative

