## Supplementary figures and images for "A phrenic-sparing cephalic reanastomosis model: Acute effects and implications"

### Figure 1

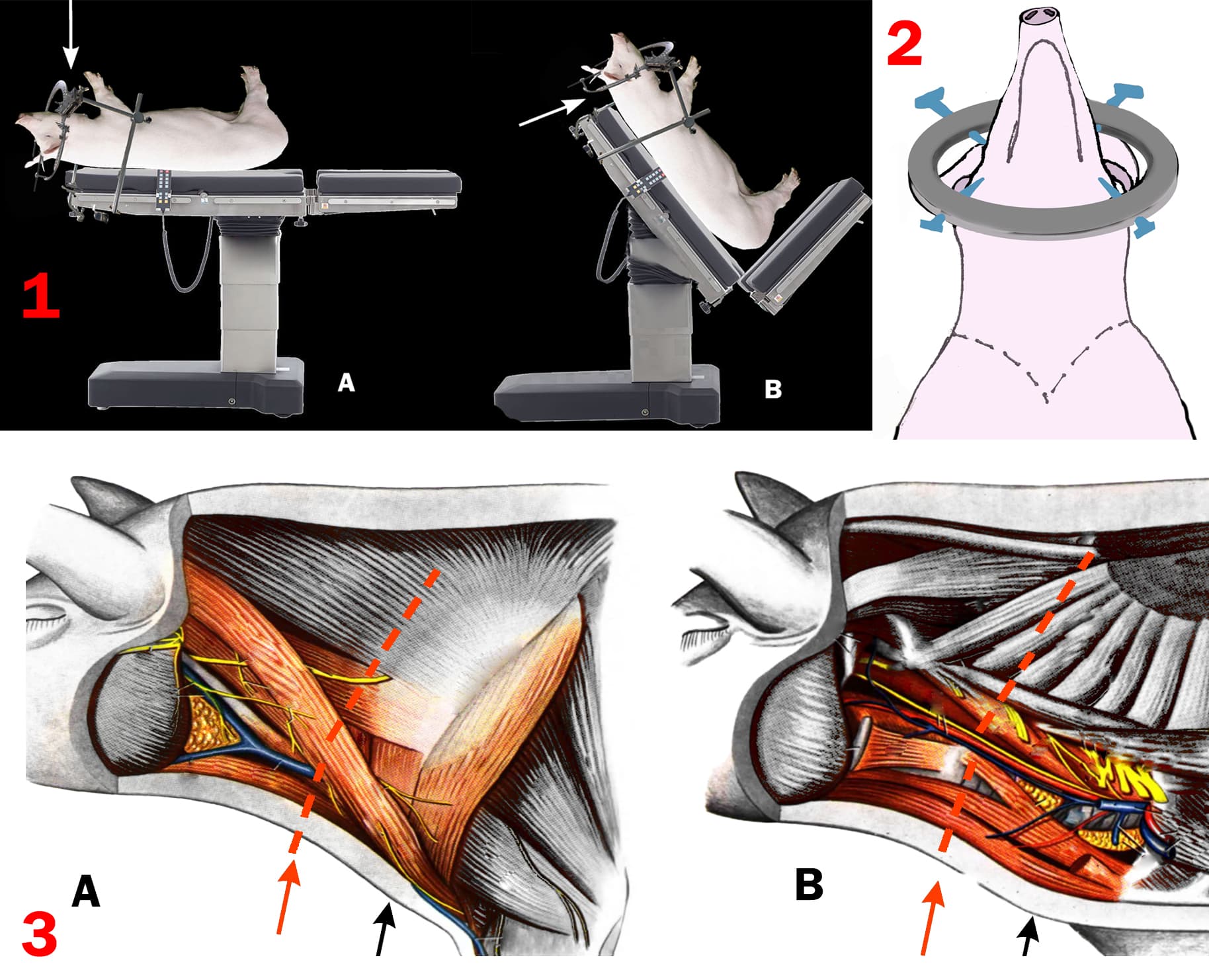

### Figure 2

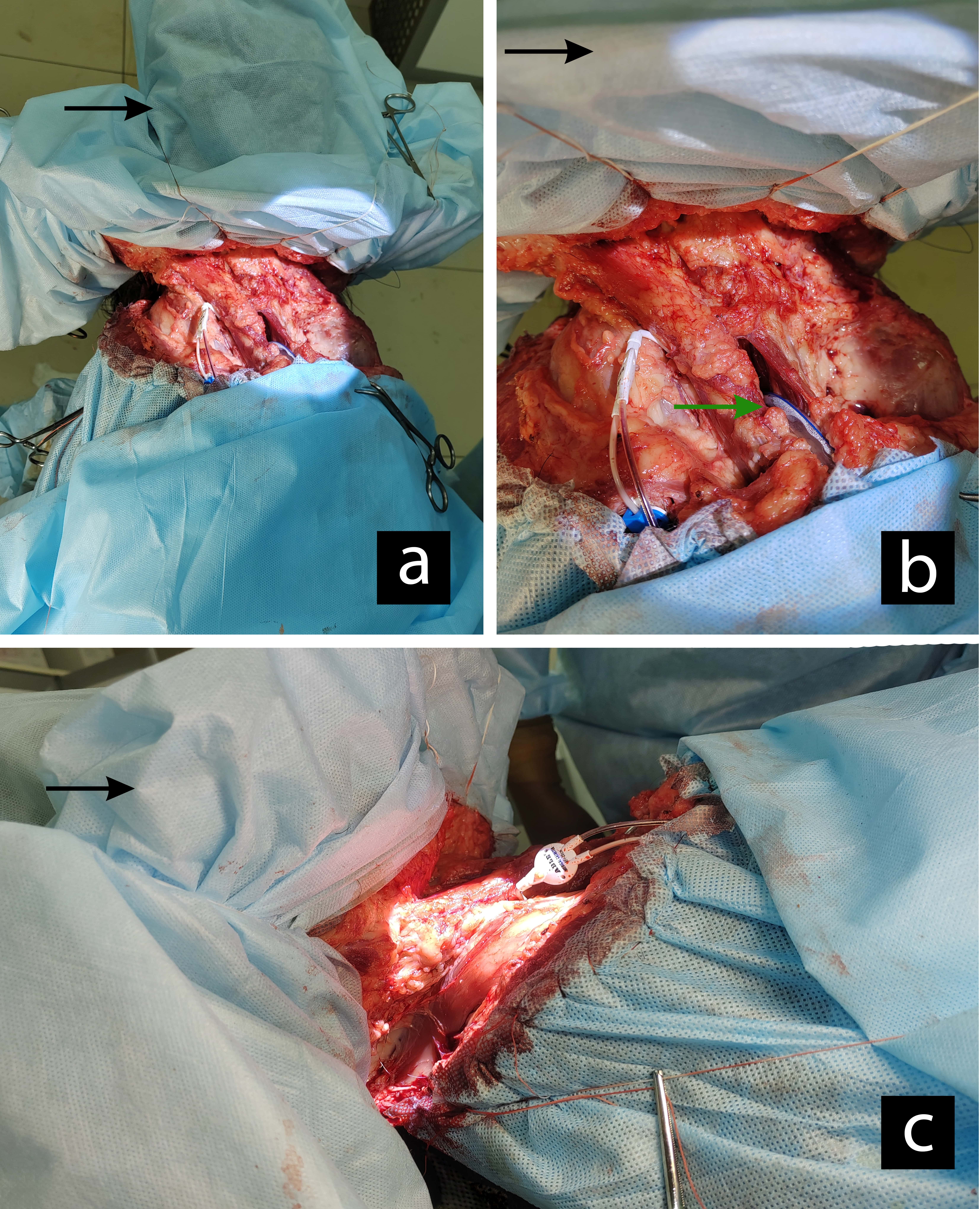

### Figure 3

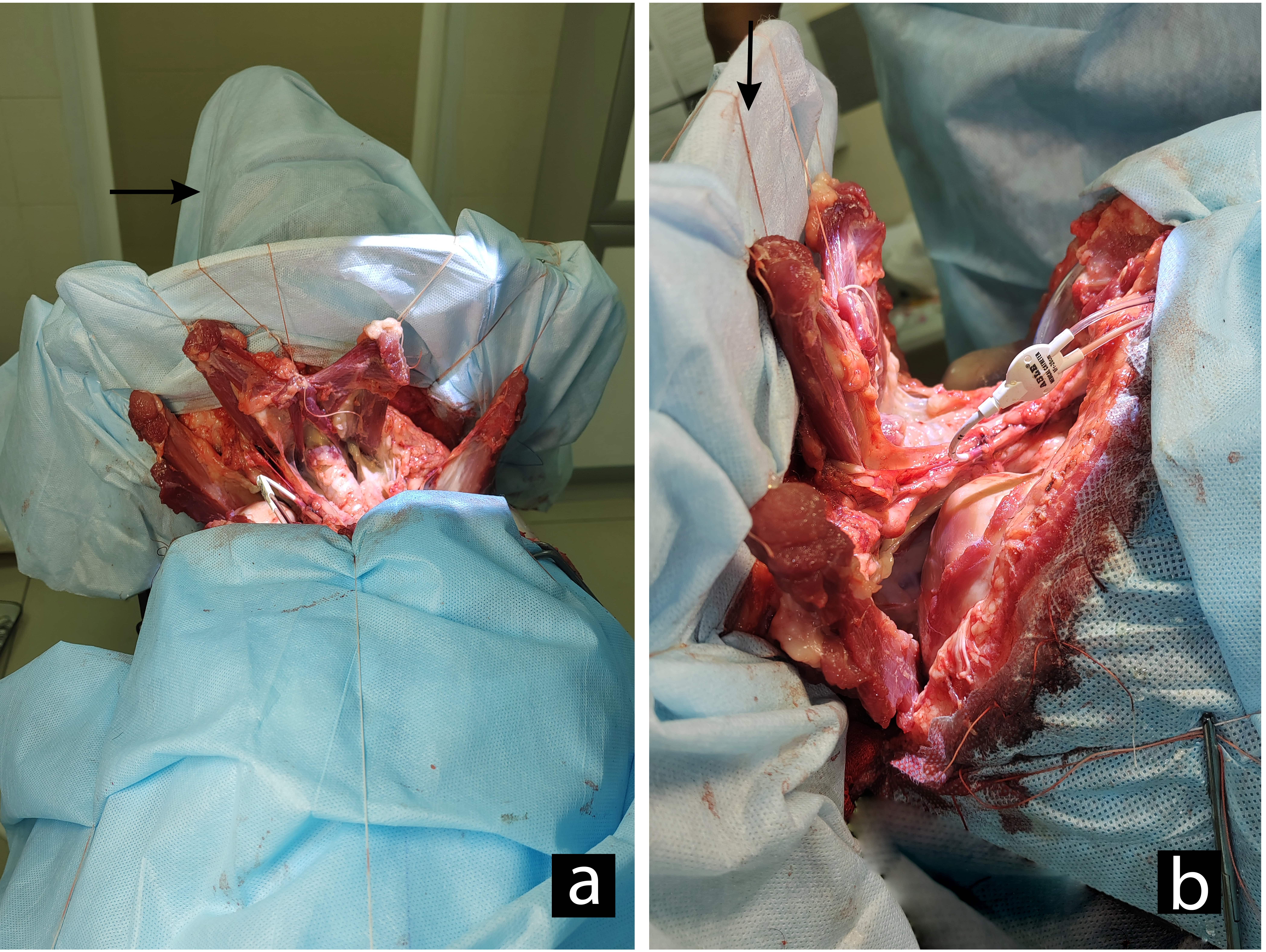

### Figure 4

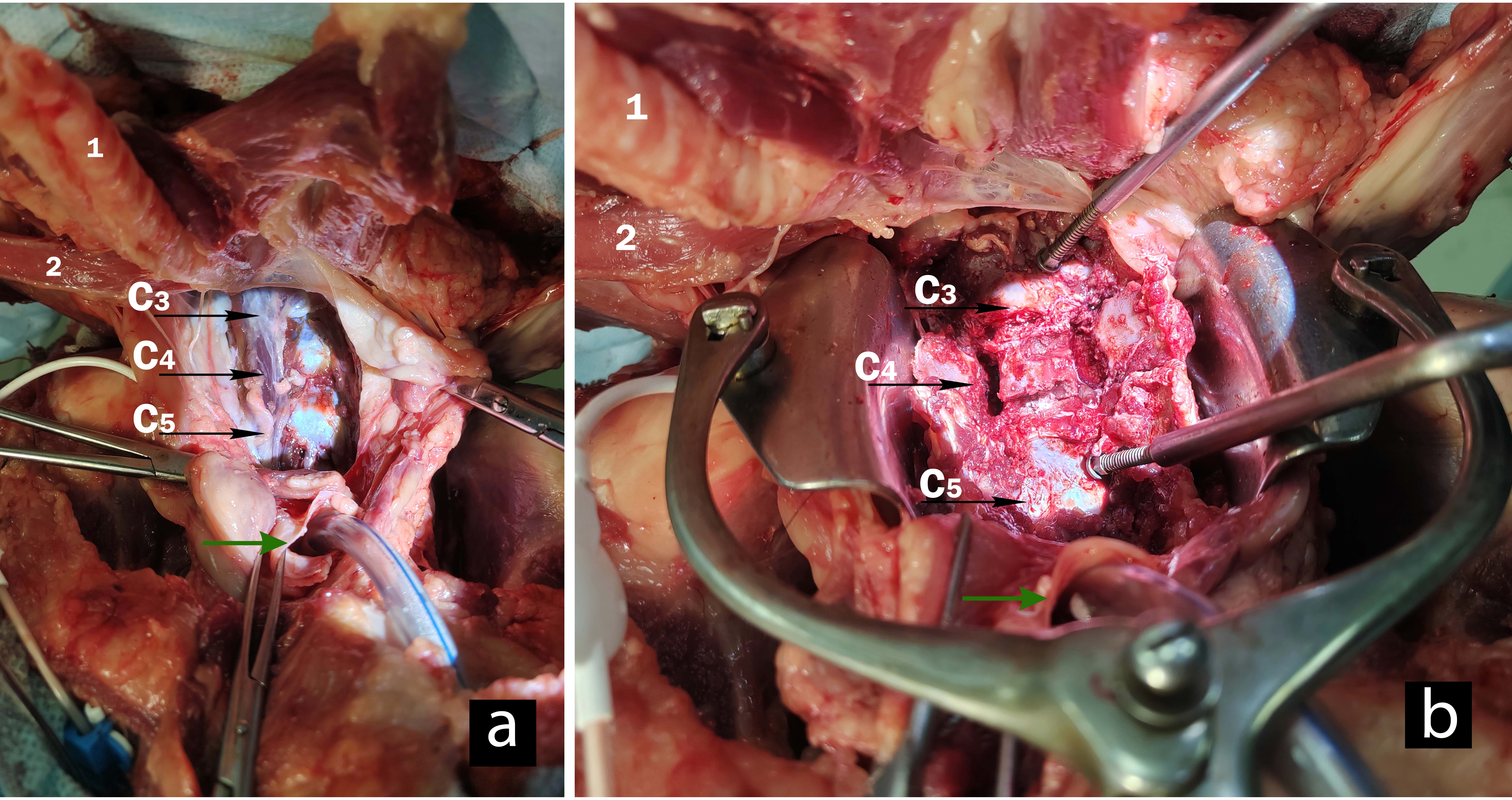

### Figure 5

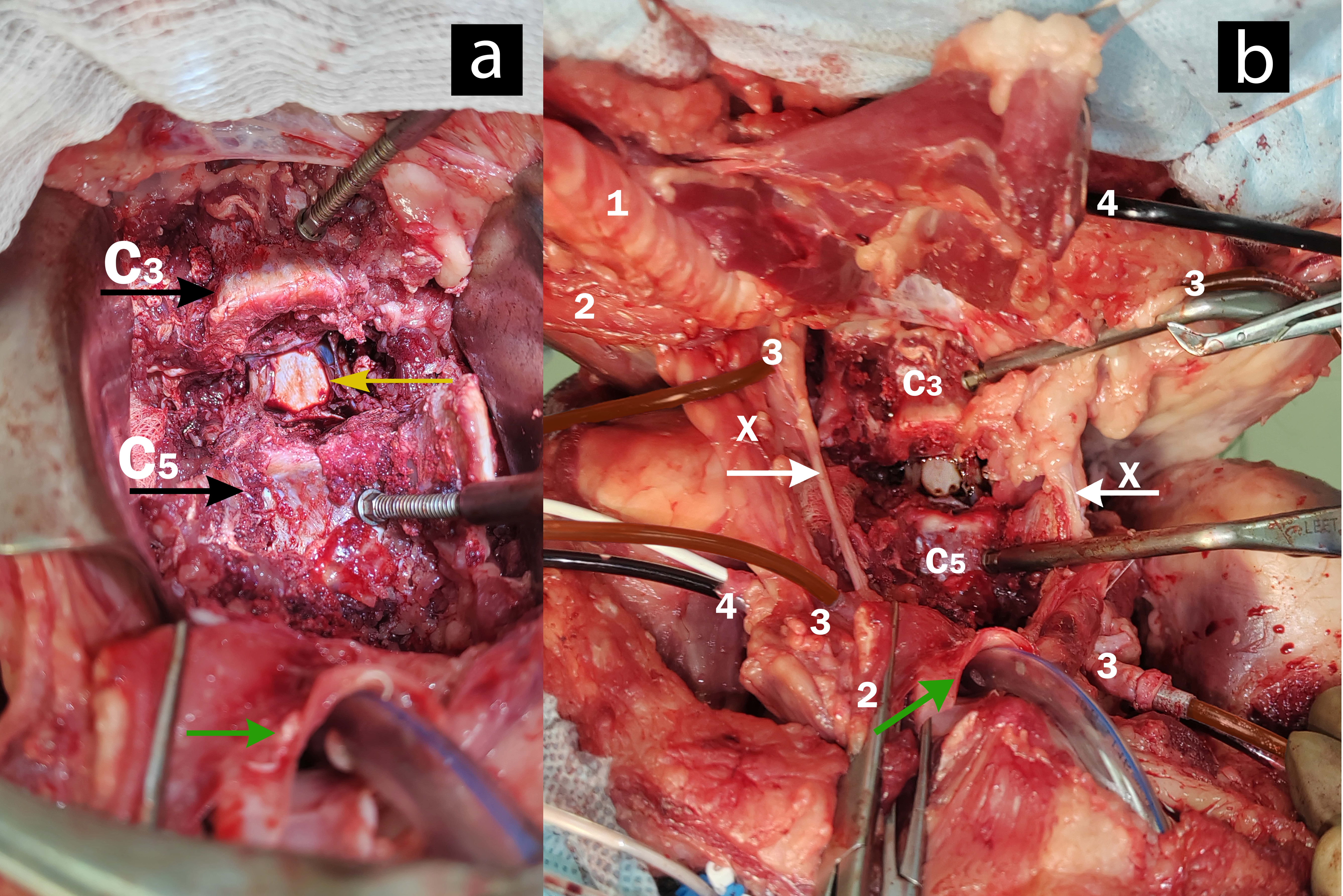

### Figure 6

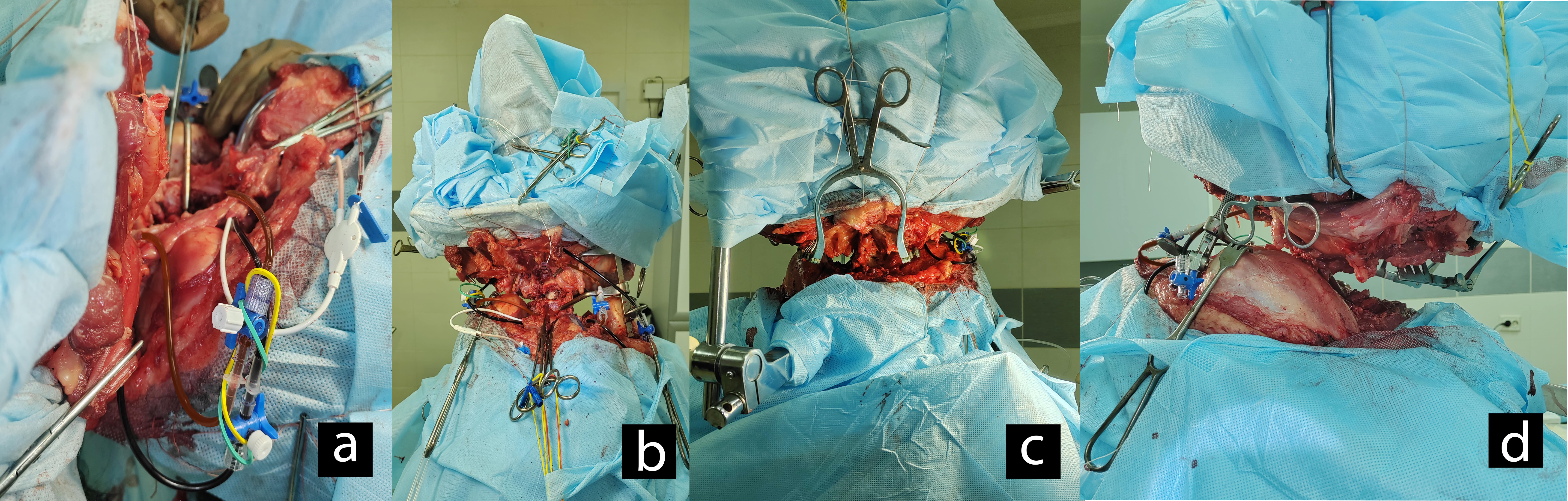

### Figure 7

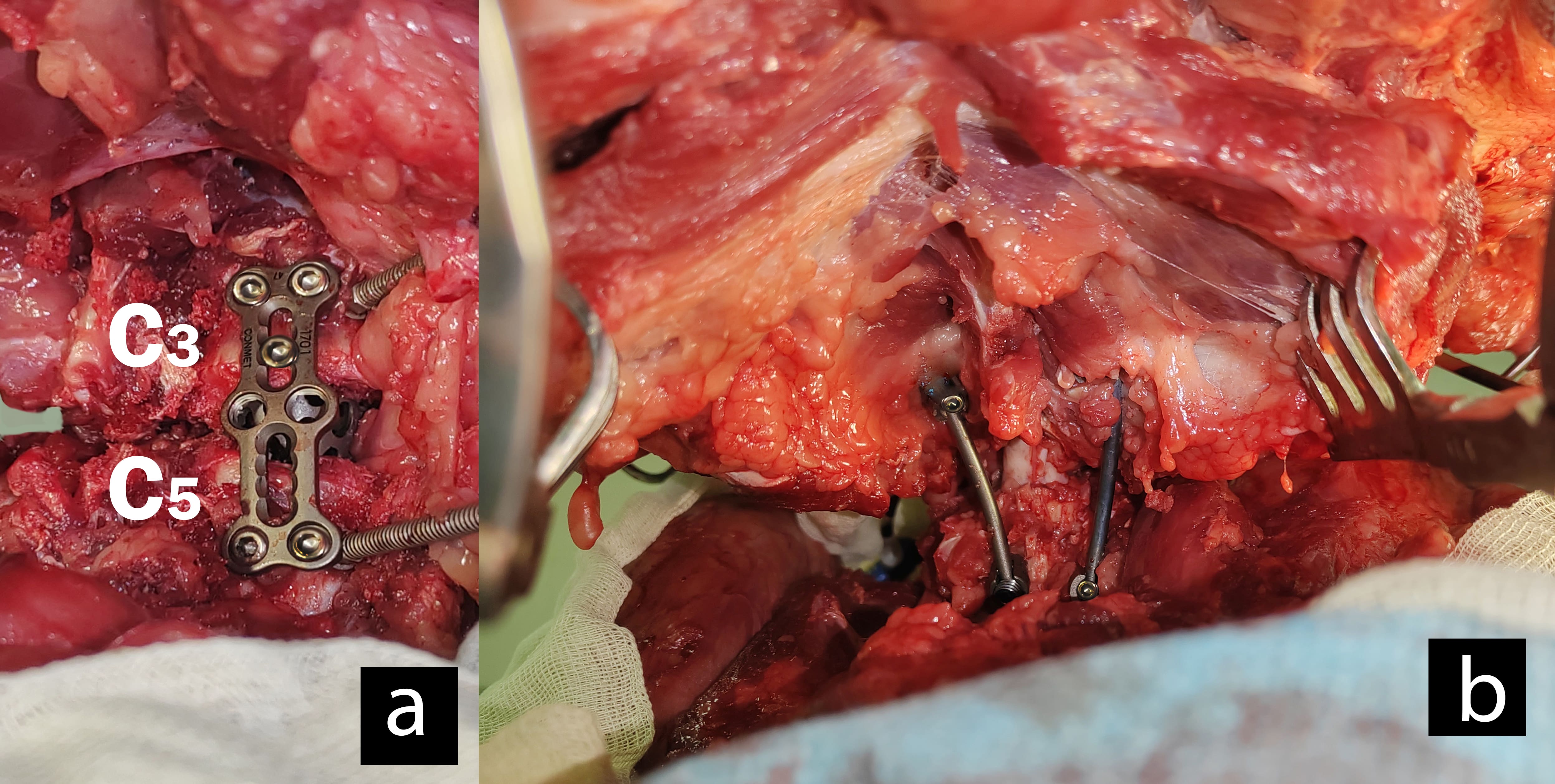

### Figure 8

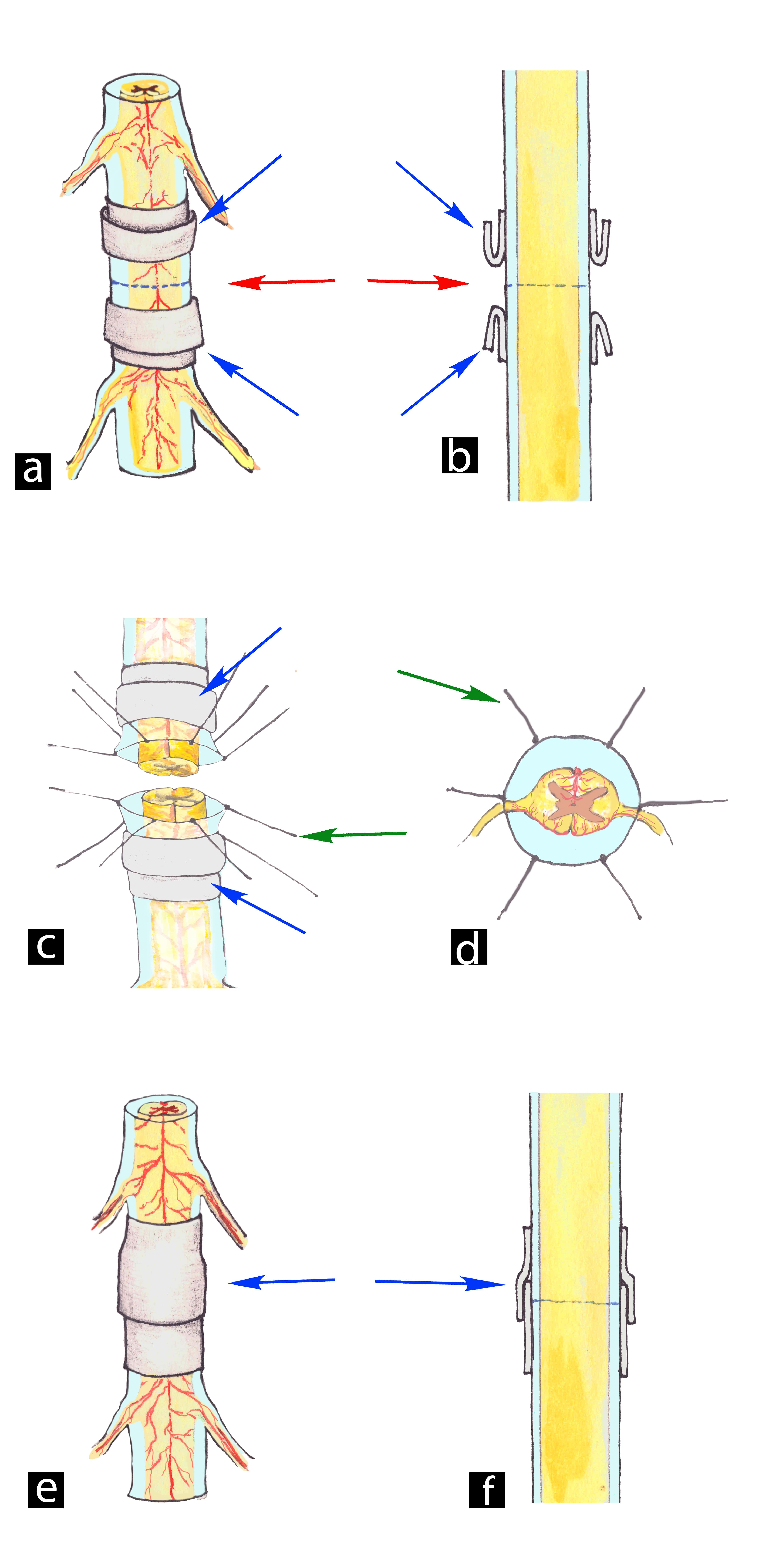

### Figure 9

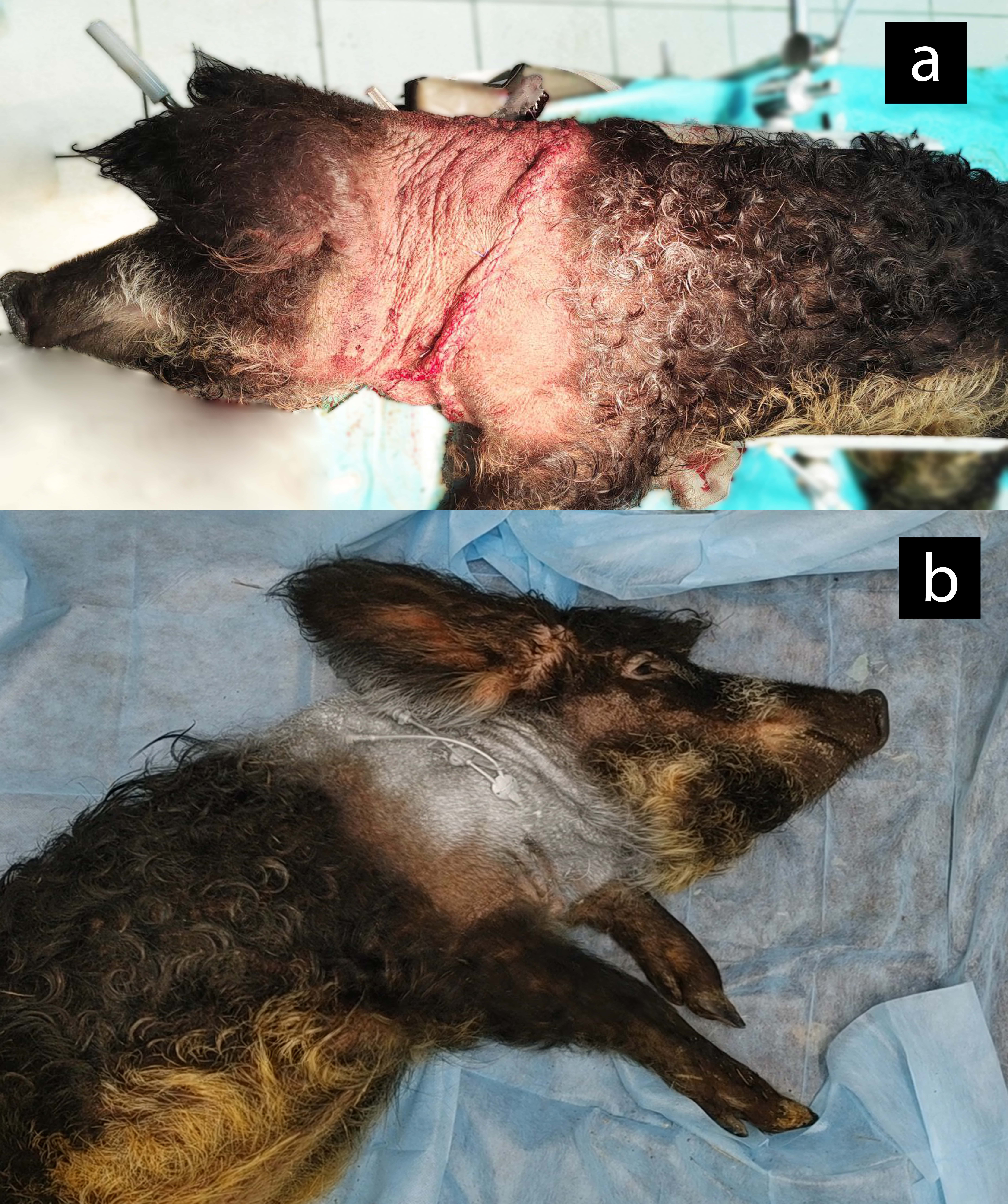
